# Partial migration in a subtropical wading bird in the Southeastern U.S.

**DOI:** 10.1101/626473

**Authors:** Simona Picardi, Peter C. Frederick, Rena R. Borkhataria, Mathieu Basille

## Abstract

The function of migration is to allow exploitation of resources whose availability is heterogeneous in space and time. Much effort has been historically directed to studying migration as a response to seasonal, predictable fluctuations in resource availability in temperate species. A deeper understanding of how different migration patterns emerge in response to different patterns of resource variation requires focusing on species inhabiting less predictable environments, especially in tropical and subtropical areas. We provide the first individual-based, quantitative description of migratory patterns in a subtropical wading bird in the Southeastern U.S., the wood stork (*Mycteria americana*). Using GPS tracking data for 64 individuals tracked between 2004 and 2017, we classified migratory behavior at the individual-year level using information theory-based model selection on non-linear models of net squared displacement. We found that the wood stork population is partially migratory, with 59% of individuals seasonally commuting between winter ranges in Florida and summer ranges elsewhere in the population range (migrants), and 28% remaining in a single area in Florida year-round (residents). Additionally, 13% of storks act as facultative migrants, migrating in some years but not in others. Comparing the distribution of residents and migrants suggests that different migratory strategies might be associated with the use of different or differently distributed resources, possibly including food supplementation from human activities. The existence of facultative migrants shows the potential for plastic change of migratory patterns. Partial migration in wood storks may be an adaptation to high heterogeneity and unpredictability of food resources. We suggest that future research should focus on wading birds as model species for the study of partial migration as an adaptation to heterogeneous and unpredictable environments, by comparing populations of the same species across different wetland systems and sympatric populations of species that differ in their resource acquisition mechanisms.

## Introduction

Migration is a widespread phenomenon across taxa, including birds, and it has the function of allowing individuals to track resources whose distribution is heterogeneous in space and time (Dingle and Drake 2007). Different forms of migration arise in response to different patterns of resource variation (Dingle 1996, Van Moorter et al. 2013). In temperate areas, where seasonality is generally repeatable, migrations take the familiar form of back-and-forth movements between ranges that are resource-rich at different times of the year (Cox 1985). However, even seemingly nomadic or irregular movements can be considered migrations if their function is to allow the exploitation of resources that do not follow seasonal fluctuations (Dingle 1996, Roshier et al. 2008, Van Moorter et al. 2013). For example, ephemeral resource outbreaks with no periodicity often lead to erratic migration (Kingsford et al. 2010). Some bird populations exhibit facultative migration when a key environmental factor that drives the availability of resources exceeds a critical threshold (Streich et al. 2006). Partial migration, when a population includes both migratory and resident individuals, often emerges when variability in the distribution of resources is paired with ecological trade-offs – such as density-dependence, the energetic cost of migration, or predator avoidance (Chapman et al. 2011). Partial and facultative migration can also be combined, when a population includes both individuals that consistently migrate and individuals that only migrate in some years (Berthold 2001, Newton 2012). Altogether, migration is a complex phenomenon encompassing a wide spectrum of behaviors which manifest as adaptations to different patterns of resource heterogeneity in space and time (Dingle and Drake 2007).

Generally, less conventional forms of migration are thought to be associated with unpredictable environments, of which wetlands are a prime example (Fletcher and Koford 2004, Niemuth et al. 2006, Sergio et al. 2011). Variation in resource distribution can happen quickly and over broad scales in wetland ecosystems (Kushlan 1986, Weller 1999). Besides within-year variability, many wetland systems are characterized by unpredictability of local conditions between years (Niemuth and Solberg 2003, Sergio et al. 2011). Accordingly, wetland-dwelling birds evolved high mobility as an adaptation to resources that pulsate unpredictably across the landscape (Haig et al. 1998, Poiani 2006). Many wading bird species (where by “wading birds” we collectively refer to Pelecaniformes, Ciconiiformes, Gruiformes, and Phoenicopteriformes; Hegemann et al. 2019) undertake large-scale movements to exploit temporary resource breakouts across the landscape (Kushlan 1981), and such movements can take many different forms and often present intra-specific differences as well (Frederick and Ogden 1997, Melvin et al. 1999, Beerens 2008).

Because they inhabit environments where resource unpredictability is brought to an extreme, wading birds seem to be a natural choice as model species to learn about the adaptive relations between migratory patterns and resource distribution. This is especially true for species inhabiting tropical and sub-tropical wetlands, where seasonality is fundamentally driven by rainfall rather than by temperature (Junk 1993). Recent literature has advocated for an increased focus on non-temperate species to deepen our understanding of migration as an adaptation to resource fluctuations in different contexts (Sekercioglu 2010). Nonetheless, few studies have explicitly quantified migration patterns of wading bird species (but see Mckilligan et al. 1993 for an example on cattle egrets, *Bubulcus ibis)* and, to our knowledge, none in non-temperate areas. In this paper, we provide a quantitative description of migratory patterns of a subtropical wading bird in the southeastern U.S., the wood stork (*Mycteria americana*).

Wood storks are distributed in the southeastern U.S. (hereafter, the Southeast), east of Mississippi and as far north as North Carolina (Coulter et al. 1999). Wood storks can travel remarkably long distances over short time frames and with low energy expenditure by soaring (Kahl 1964, Ogden et al. 1978). This is an adaptation to high heterogeneity and unpredictability of food resources, which are an important driver of wood stork population responses (Frederick and Ogden 2001, Gawlik 2002, Herring 2008). Wood storks are tactile foragers that feed almost exclusively on fish (Kahl 1964, Ogden et al. 1976, Kushlan 1986). For them to forage efficiently, prey need to be highly concentrated (Kahl 1964, Kushlan 1986, Gawlik 2002). As a result of local differences in hydrological dynamics, high fish concentrations occur at different times and in different locations within wetland systems in the wood stork range, and they are generally ephemeral (Loftus and Eklund 1994, Frederick et al. 2009, Botson et al. 2016). For example, in the Florida Everglades, where historically most wood stork nesting activities occurred in the U.S. (Frederick and Ogden 1997), high water levels promote the growth of fish populations during the rainy season (DeAngelis et al. 2010, Botson et al. 2016). Then, as the water recedes in the dry season, retention of pockets of water in shallow depressions across the landscape concentrates fish, making them available for birds (Kahl 1964, Kushlan 1986, Frederick et al. 2009). The result is a spatio-temporally heterogeneous mosaic of foraging habitat, where food availability changes rapidly through time and space due to the interaction of hydrology and topography (Chick et al. 2004, Ruetz et al. 2005, DeAngelis et al. 2005). Other wetland systems in the Southeast may present different phenologies and mechanisms of food concentration, but their hydrological dynamics are also largely influenced by rainfall patterns, affecting the distribution of resources (Snodgrass et al. 1996, Baber et al. 2002).

Wood stork movements reflect patterns of resource availability at fine spatio-temporal scales. For example, during the breeding season, wood storks move long distances from breeding colonies to foraging grounds to accommodate shifting resource availability patterns (Kahl 1964, Ogden 1986, Bryan and Coulter 1987). At a broad spatio-temporal scale, the annual range of wood storks includes wetlands located in different states that are subject to different, sometimes asynchronous, and usually unpredictable rainfall patterns (Frederick et al. 2009). For example, southern Florida is a winter dry, summer wet monsoonal system, while much of the rest of the southeast gets most of its rainfall in winter and dries during summer months. Because their range includes wetland systems subject to different climatic regimes, local conditions within seasonal ranges used by wood storks are characterized by high year-to-year unpredictability as well (Gawlik 2002, Frederick et al. 2009). Landscape-scale movements of wood storks might respond to heterogeneity in food resources at this scale similarly to how fine scale movements reflect heterogeneity in food availability patterns within seasonal ranges.

Landscape-scale movements of wood storks remain poorly understood. Previous literature reported large scale movements of wood storks between different parts of their U.S. range in different seasons, but defined the species as “not a true migrant” (Coulter et al. 1999). Indication that movements between different areas within the range are repeated year after year, thus presenting typical migration features, was provided by a study on juveniles (Hylton 2004). Other tracking studies have described fine-scale movements (Borkhataria et al. 2013) or dispersal (Bryan et al. 2008, Picardi et al. 2018), but not migration. Overall, we lack a formal understanding of the migratory status of the wood stork population.

The objective of this study was to address the outlined knowledge gaps by providing an individual-based, quantitative description of wood stork migratory patterns in the Southeast. By leveraging a large, long-term GPS-tracking database, we quantified migratory behavior of a large number of individuals over 14 years and evaluated behavioral consistency across years at the individual level. We determined correlations between migratory behavior and life-history traits including age and sex. We mapped seasonal distribution patterns of wood storks as a result of migration patterns and quantified individual site-fidelity. Our results contribute to our general understanding of the adaptive significance of partial migration in unpredictable environments by providing the first individual-based documentation of migration patterns in a subtropical wading bird.

## Methods

### Wood stork captures and data collection

We used GPS telemetry to track wood stork individual movements throughout the population range (centroid 28.8967°N, 81.3310°E) between 2004 and 2017. A total of 133 wood storks were captured at 11 sites throughout the population range (Figure 1) either by hand (in the case of juveniles) or using rocket nets. Captured storks were hooded to reduce stress during handling and equipped with solar-powered GPS transmitters (Microwave Telemetry, Columbia, MD), which are not limited by battery life. The transmitters were programmed to record a location every 1 (*n* = 112) or 2 hours (*n* = 21).

**Figure 1.**
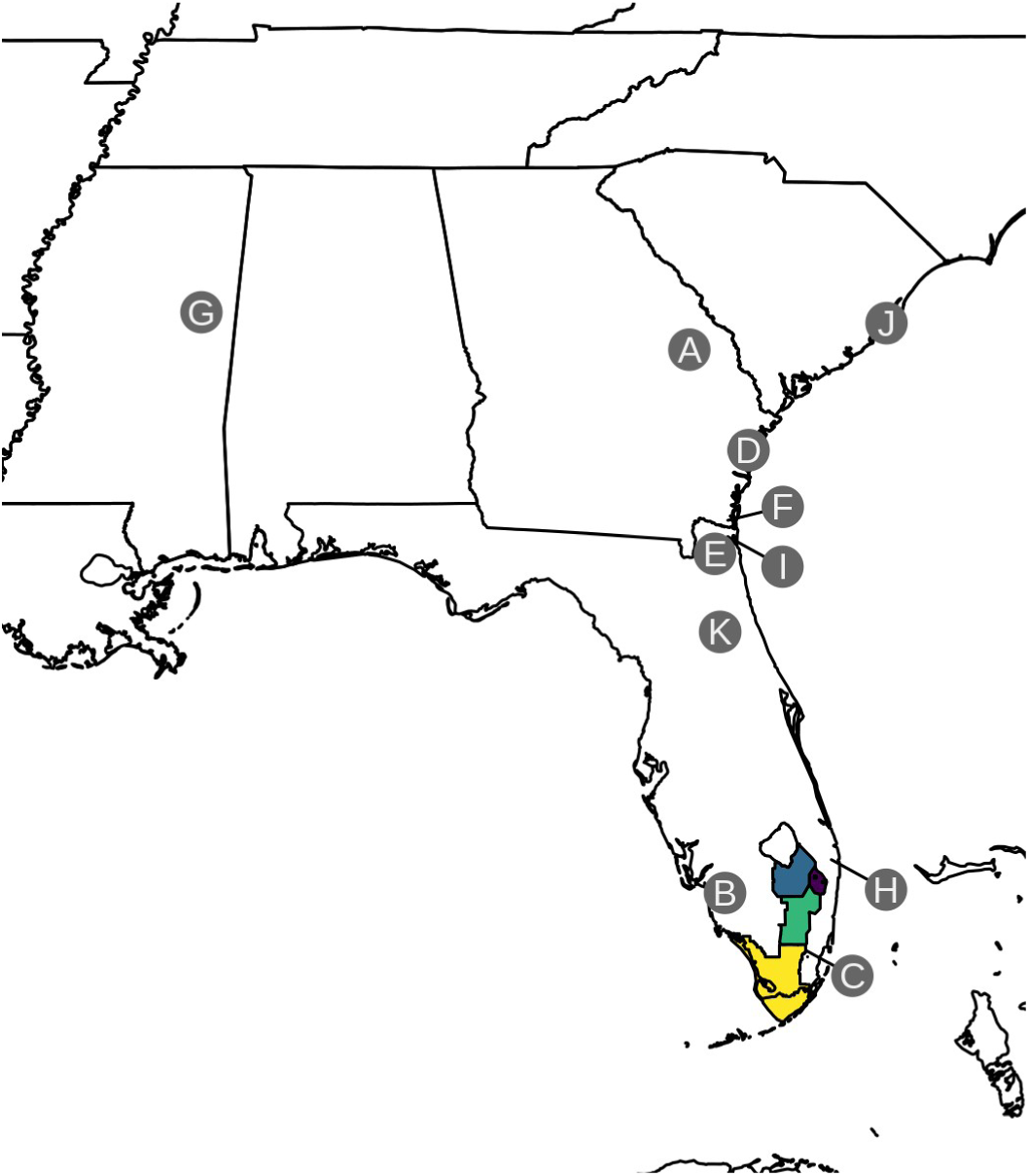
Map of the study area, located within the breeding range of the southeastern U.S. wood stork population. The letters indicate capture sites. We indicate in parentheses the number of wood storks captured at each site: A = Chew Mill Pond (*n* = 2), B = Corkscrew Swamp Sanctuary (*n* = 4), C = Everglades National Park (*n* = 9), D = Harris Neck National Wildlife Refuge (*n* = 10), E = Jacksonville Zoo (*n* = 9), F = Kings Bay Naval Base (*n* = 1), G = Noxubee National Wildlife Refuge (*n* = 2), H = Palm Beach Solid Waste Authority (*n* = 14), I = St. Mary’s (*n* = 1), J = Washo Preserve (*n* = 9), K = Welaka Fish Hatchery (*n* = 3). The colored polygons depict the boundaries of relevant management units within the Everglades watershed. Blue polygon = Everglades Agricultural Area, Purple polygon = Loxahatchee National Wildlife Refuge, Green polygon = Water Conservation Areas 2 and 3, Yellow polygon = Everglades National Park.

### Operational definitions

We operationally defined migration as a round-trip between ranges that were spatially separated and used at different times during the year – thus implying return to the initial range. We defined migratory choice as a binary variable at the year level, namely whether an individual migrated or not in a given year. We then combined migratory choices for an individual in different years to assess multi-year migratory strategies. Thus, we defined migratory strategy as the history of yearly migratory choices of an individual. For example, an individual whose migratory choice is migration every year adopts a pure migrant strategy, or an individual whose migratory choice is different in different years shows a facultative migrant strategy.

### Classification of migratory behavior

We used GPS tracking data to investigate individual migratory choices based on net squared displacement (NSD), i.e. the squared linear distance between any point along a movement trajectory and an arbitrarily chosen starting point (Kareiva and Shigesada 1983, Calenge et al. 2009). This metric provides an intuitive measure of how far an individual is from a reference point in space at any time (Kareiva and Shigesada 1983, Calenge et al. 2009). To classify wood stork migratory behavior at the yearly scale, we used a modeling approach adapted from a method first introduced by Bunnefeld et al. (2011) and later improved by Spitz et al. (2017). The approach consists of fitting a set of non-linear models to yearly individual NSD time series and selecting the one that best fits the data using AIC. For the purpose of our study, following our binary definition of migratory choice, we took two possible models into consideration: a migrant model and a resident model (Figure 2; see Spitz et al. 2017 for details on model specification). In the migrant model, the yearly time series of NSD follows a double sigmoid curve, indicating initial residency in one range (initial low NSD phase), displacement to a second (high NSD phase), and subsequent return to the initial range (final low NSD phase; Figure 2). The departure range is whichever range an individual was located in at the arbitrarily chosen reference time that marks the start of the trajectory. The resident model is represented by a horizontal asymptotic curve, indicating permanence in a single range after an initial phase of increase of NSD until settlement around a constant value (Figure 2). We applied model selection based on Akaike’s Information Criterion (AIC) differences on these two competing models to classify wood stork annual trajectories and determine migratory choice. The analysis was performed in R (R Core Team 2018) using functions implemented in the migrateR package (Spitz et al. 2017).

**Figure 2.**
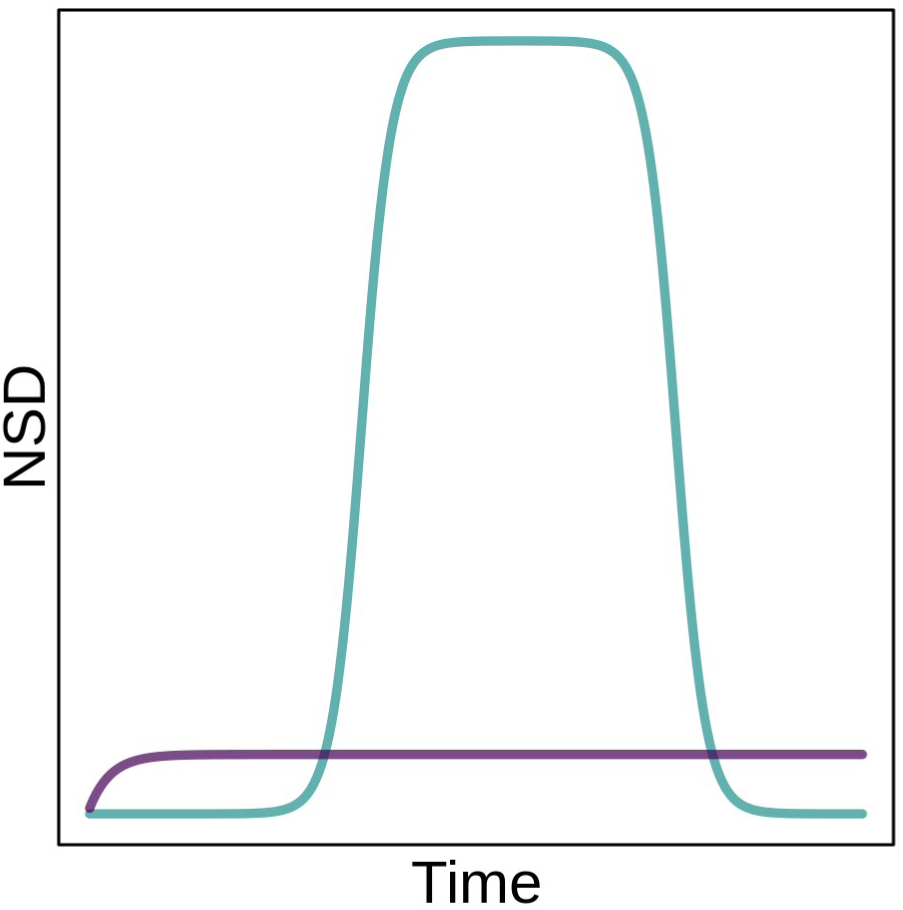
Conceptual illustration of non-linear models of Net Squared Displacement used to classify wood stork migratory behavior at the year scale. Resident model in purple, migrant model in blue. Figure adapted from Bunnefeld et al. (2011) and Spitz et al. (2017).

### Data preparation

After visual exploration of the trajectories, we divided the tracking data into yearly individual trajectories starting on January 15^th^, to minimize the probability of the starting point falling within a migration (following recommendations in Spitz et al. 2017). We used the R packages adehabitatLT (Calenge 2006) and rpostgisLT (Dukai et al. 2016) for data processing and exploration, respectively. We screened the resulting yearly individual trajectories to assess whether they included sufficient data for model fitting. In order to ensure detection of migrations, we set the minimum data requirements for an individual-year to at least 15 locations every 4 months (Jan-Apr, May-Aug, Sep-Dec). The resulting dataset consisted of 212 individual-years from 66 storks, of which 20 had a single individual-year and 46 had multiple individual-years (range = 2-10, mean = 3.96 ± 1.73 SD).

### A-priori model constraints

Following recommendations in Spitz et al. (2017), we enforced a-priori constraints in the model parameters to satisfy the following quantitative characterization of migration: for an individual to be considered a migrant on a given year, it has to spend at least 60 days in a range at least 260 km away from the departure range. The chosen spatial threshold corresponds to double the maximum distance documented for wood stork trips from the colony to foraging grounds (130 km; Kahl 1964, Ogden et al. 1978), which is presumably a distance that storks are able to cover within their everyday movements. Thus, this value seems appropriate to discriminate between the scale of within-versus between-ranges movements. Temporal fluctuations of resource availability usually occur with seasonal (i.e. multiple months) periodicity at a broad spatial scale in the wood stork population range, which is expected to reflect in the emergence of migration as a seasonal phenomenon. The function of repeatedly tracking resource availability over broad spatial and temporal scales is what distinguishes migration from other types of movements which were not the focus of this study. Thus, the chosen temporal threshold of approximately 2 months has the purpose of preventing brief but spatially broad excursion movements, which are functionally different from migration, from being misclassified as migrations. We performed a sensitivity analysis on the use of different constraint values (see Appendix A) and found both the chosen spatial and temporal thresholds to be conservative, since classification results were robust to the use of a broad range of values around the chosen one, within a range of biologically meaningful values.

### Stepwise specification of starting parameter values

In addition to specifying constraints for two of the model parameters as described above, we ensured model convergence on all trajectories by progressively specifying different starting values for model parameters, following recommendations in Spitz et al. (2017). These include, for the migrant model, the midpoint of the departing migratory movement, the duration of the migratory movement, the permanence time in the arrival range, and the distance between seasonal ranges (Spitz et al. 2017). For the resident model, parameters include the average NSD of the resident range and the rate of the initial NSD increase (Spitz et al. 2017). Stepwise manual specification of starting parameter values facilitates parameter optimization, helping to overcome commonly encountered convergence issues due to the use of a single set of starting values for all trajectories in a sample (Spitz et al. 2017). All models converged after 21 iterations with a different set of starting parameters (see Appendix B).

### Post-hoc model evaluation

Following recommendations in Spitz et al. (2017), we visually inspected results of model fitting as a *post hoc* evaluation. While the minimum data requirements we chose were adequate in most cases (200 individual-years), for 12 individual-years the placement of the 15+ locations within the first or third quadrimester did not allow for an unequivocal classification (see Appendix C). The most common issue was failure to classify seemingly migratory individual-years as migrants because of insufficient temporal cover of the data (*n* = 8): while the first range shift was identified, the return movement from the second range and the subsequent residency back in the first range were not captured in the data, resulting in a poor fit of both the migrant and the resident model. Conversely, 4 individual-years that did not seem to exhibit migratory behavior, and were thus classified as residents, had similar limitations in terms of temporal cover of data that did not allow for reliable classification: we cannot exclude that migrations were not observed because they simply happened outside of the tracking period. Therefore, we discarded these 12 individual-years from further analyses. Out of the remaining 200 individual-years, 2 consecutive years for one migrant individual were erroneously classified as residents because the individual migrated over the start of the new tracking year, resulting in the migratory movement being split into two. For this individual only, we repeated model fitting and selection after splitting the yearly trajectories on December 1^st^ instead of January 15^th^. Finally, by visual comparison between model output and mapped trajectories, we identified 2 individual-years that were classified as migrations as controversial cases. These individuals continuously performed movements at a broader spatial scale than other resident individuals in the sample, but without a clear pattern of seasonal periodicity or spatial separation. Because these movements did not fit the chosen definition of migration, we manually assigned these individual-years to the resident category.

### Seasonal distributions, range fidelity, and migratory consistency

The output of the NSD models provided estimates for key migratory parameters, including the time of migration start and end where applicable. Based on these, we subsetted individual tracking datasets into residency and migration phases. For migrant individuals, we computed seasonal home ranges (winter and summer) using locations during residency phases only (i.e., excluding locations during migration trips). For resident individuals, we computed both year-round home ranges, and seasonal home ranges using locations included between the mean spring and fall migration dates observed in the population. All home ranges were computed using the kernel density estimation method, extracting the 90% density isopleth of the utilization distribution, as recommended by Börger et al. (2006), using the R package adehabitatHR (Calenge 2006). We used linear mixed models to assess differences in home range size between migrants and residents in each season while taking individual variation into account. We log-transformed home range size before fitting the model. We fit an interaction between season and migratory behavior and added the individual identity as a random effect. We evaluated model predictions at the fixed effects level to assess differences between migrants and residents in different seasons. We evaluated model fit using the pseudo-R^2^ method of Nakagawa and Schielzeth (2013). For individuals that were tracked for multiple years (*n* = 46), we investigated seasonal range fidelity using home range overlap, with 1 representing perfect overlap and 0 representing disjunct ranges. For summer and winter separately, we computed the percent area of overlap between all pairwise combinations of ranges of each individual, and averaged them in a synthetic value of individual site fidelity. Finally, we quantified migration strategies at the individual level by modeling migratory choices of an individual in different years as a binomial process. Using the binomial likelihood, we computed maximum likelihood estimates of individual migration probability along with 95% confidence intervals. Values reported in the Results are means ± SD.

## Results

### Migratory choices and strategies

The final classification of wood stork migratory choices consisted of 200 individual-years from 64 individuals (15 captured as juveniles of unknown sex, 25 adult females, and 24 adult males), of which 121 were migrations (∼60%) and 79 (∼40%) residencies. The maximum likelihood estimates of migration probability, which describe individual migratory strategies, were 1 for 36 individuals (of which 27 tracked for multiple years), 0 for 22 (of which 13 tracked for multiple years), and between 0 and 1 for 6 (all tracked for multiple years; Figure 3). Confidence intervals around the estimates of migration probability were large due to the limited number of tracking years (range 1-10). Among the individuals tracked for multiple years, 40 showed consistent migratory choices across years, thus adopting a pure migrant (∼59%) or pure resident strategy (∼28%), while 6 (∼13%) showed variable migratory choices across years. Among pure migrants, 15 were male and 16 female (5 were of unknown sex); among pure residents, 6 were male and 7 female (8 were of unknown sex); among individuals with variable migratory choices, 3 were male and 2 female (2 were of unknown sex). Among the 15 storks that were captured as juveniles, 8 were tracked into subsequent years as subadults (*n* = 8) or adults (*n* = 2). Of these, 6 exhibited consistent behavior across years (3 pure migrants and 3 pure residents), while 2 showed variable migratory choices in different years. Overall, we found no correlation between individual migratory choices and age or sex.

**Figure 3.**
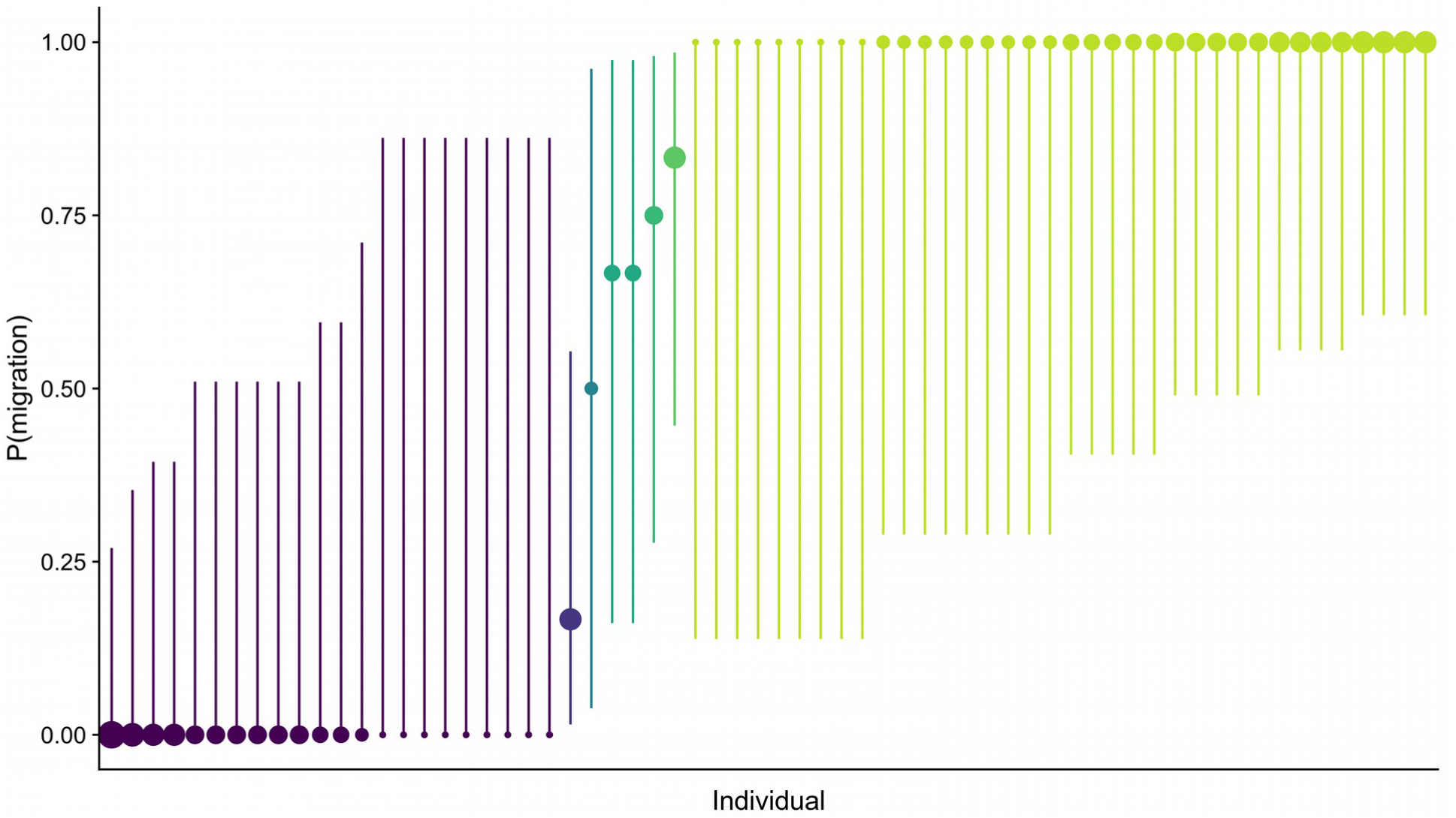
Maximum likelihood estimates of individual migration probabilities with 95% confidence intervals. The size of the points is proportional to the number of tracking years for each individual.

### Migration routes and timing

The mean departure dates were May 7^th^ and October 2^nd^ for spring (*n* = 121) and fall migrations (*n* = 121), respectively. The distribution of migration departure dates was bimodal in spring, with a peak in late March and one in June (Figure 7A), while departure dates in fall showed an early surge followed by a single peak in mid-October (Figure 7A). Storks followed two general migration routes along the east and west coastline of Florida, with the east one used more in spring and the west in fall (Figure 7B).

### Seasonal ranges and population distribution

The overall population distribution (including both migrants and residents) was highly dispersed throughout the Southeast in summer, while highly concentrated in south Florida in winter (Figure 4). The year-round ranges of resident individuals were concentrated in a few hotspots in southeast Florida and in the Jacksonville area (Figure 5). The interaction between season and migratory behavior significantly affected home range size (*p* < 0.05). The marginal R^2^ was 0.07 and the conditional R^2^ was 0.44, suggesting that individual variability explained most of the variance rather than the fixed effects. Migrants, but not residents, showed larger seasonal ranges in winter than in summer (Figure 6). Migrant winter ranges were larger than both summer and winter ranges of residents (Figure 6). Wood storks exhibited moderate range fidelity (migrants = 0.51 ± 0.37 and residents = 0.62 ± 0.38 in winter, migrants = 0.51 ± 0.43 and residents = 0.61 ± 0.41 in summer).

**Figure 4.**
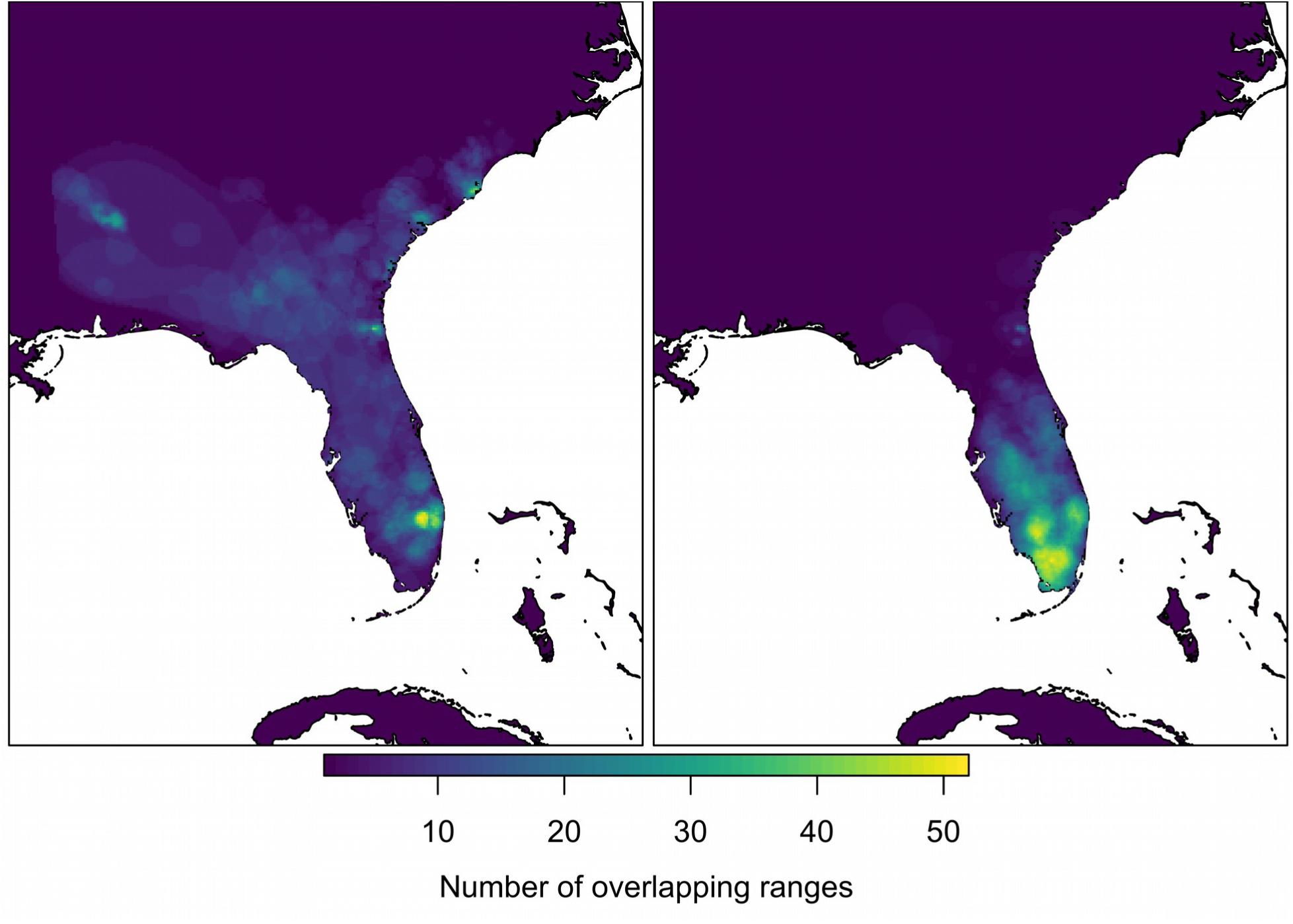
Heat map of wood stork population distribution in summer (left panel) and winter (right panel). Seasonal ranges of both migrant and resident individuals are included. Home ranges used in different years are overlaid.

**Figure 5.**
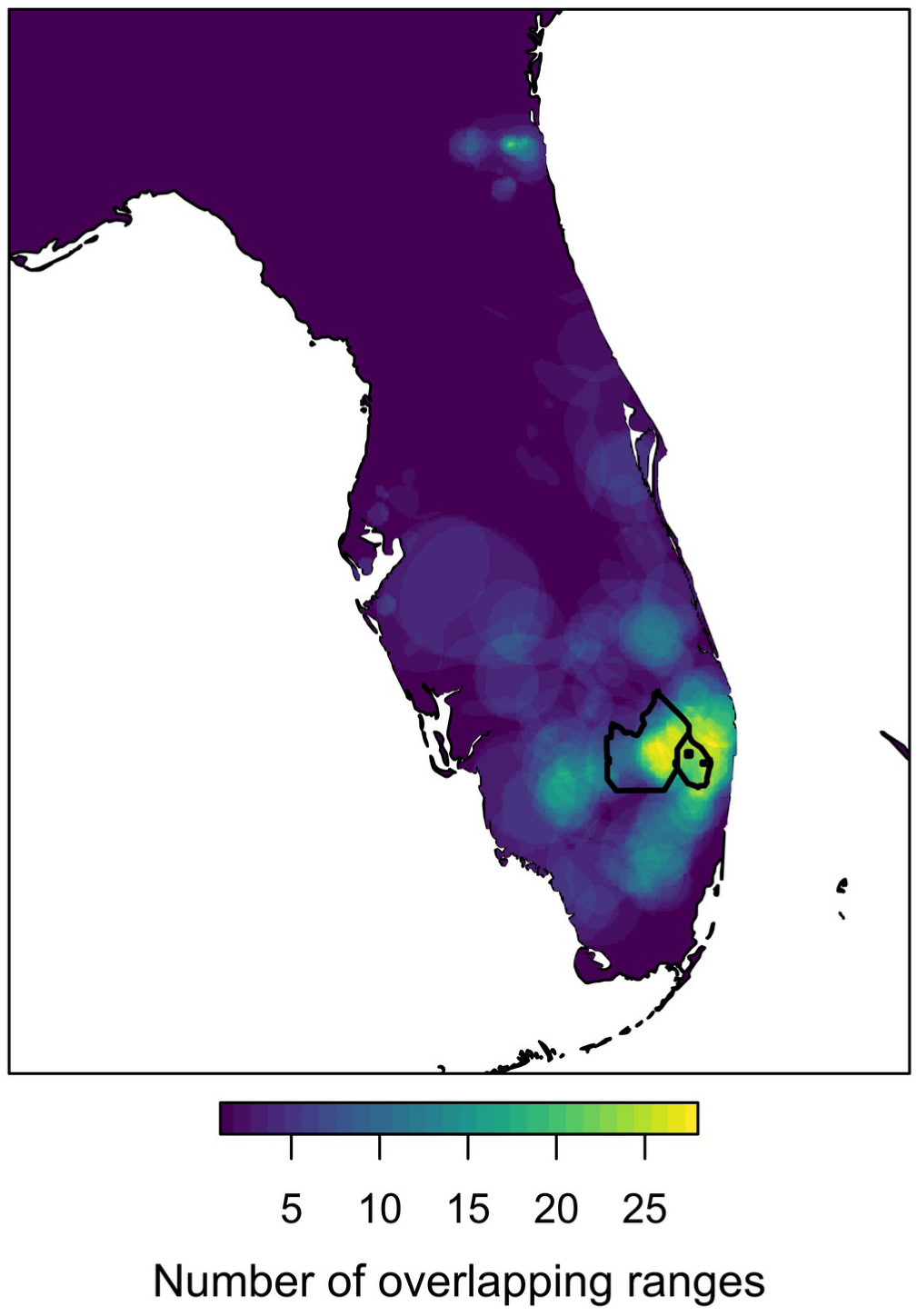
Heat map of year-round distribution of resident wood storks. Yearly ranges used by resident individuals in different years are overlaid. The black outlines depict the boundaries of Everglades Agricultural Area (left polygon) and Loxahatchee National Wildlife Refuge (right polygon).

**Figure 6.**
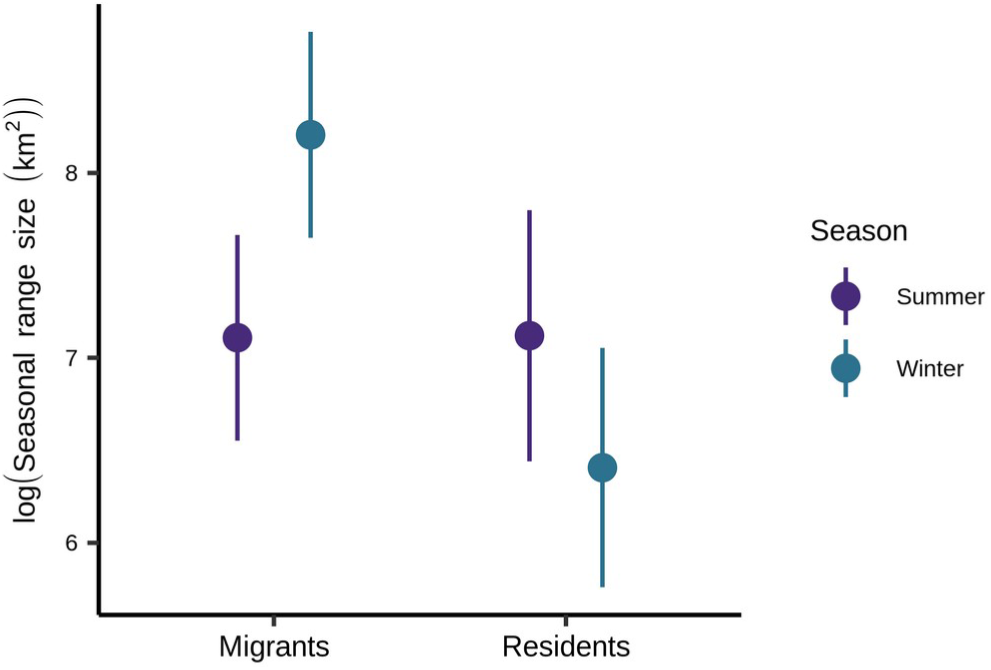
Model predictions for seasonal range size of migrants (left) and residents (right) in summer (purple) and winter (blue).

**Figure 7.**
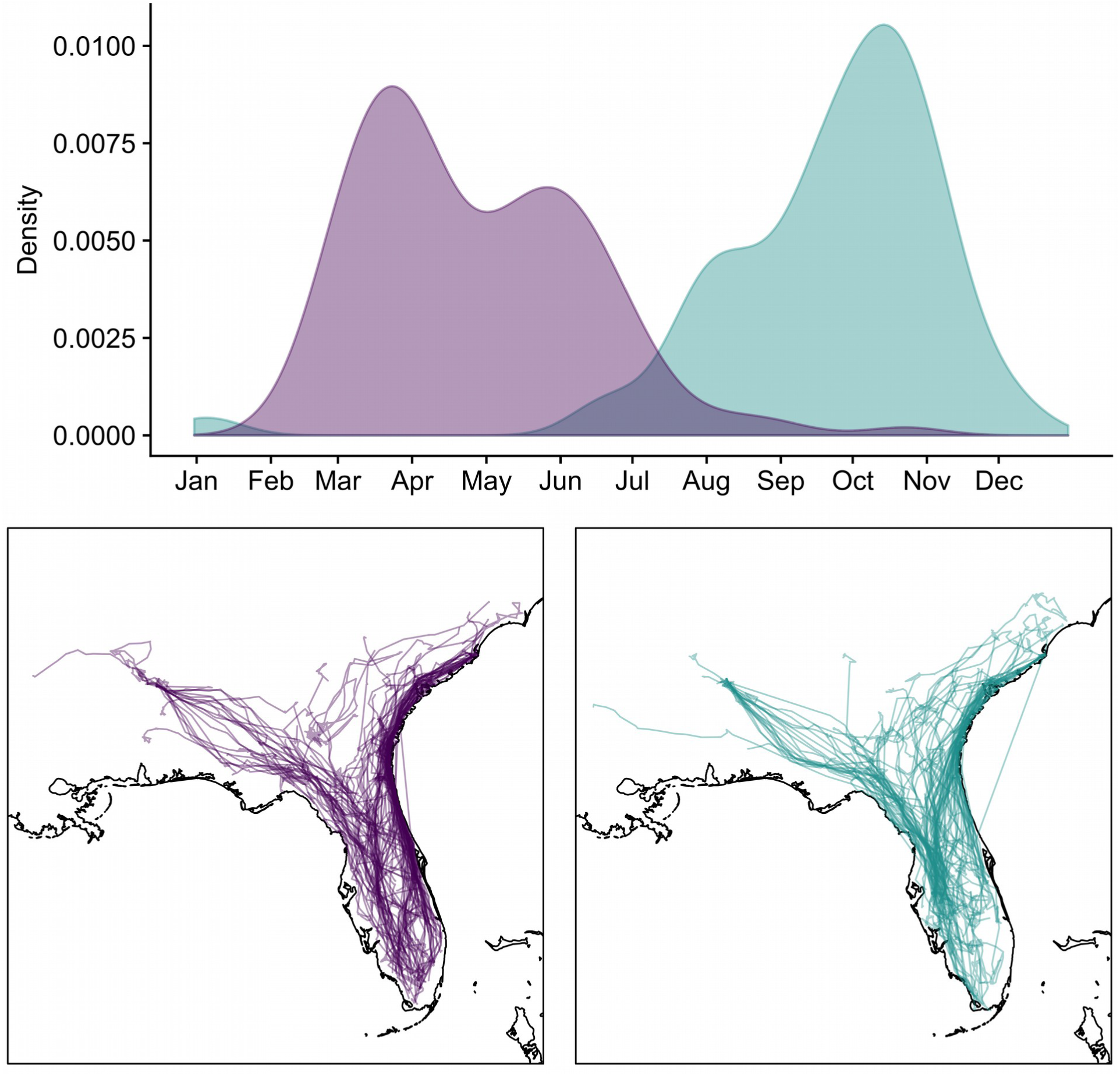
Wood stork migration routes and timing. (**A**) Frequency distribution of departure times for spring (purple) and fall (blue) migration. (**B**) Routes of migration for spring (purple, left panel) and fall (blue, right panel).

## Discussion

We provided an individual-based quantitative description of migratory patterns in a subtropical wading bird, the wood stork, in the southeastern U.S. Our findings revealed that the population is partially migratory, with a group of individuals that seasonally commute between spatially distinct ranges and others that remain resident in the same area year-round. Migration and residency appeared to be alternative choices adopted by different individuals, but less frequently by the same individuals in different years. Thus, the population exhibits a combination of partial and facultative migration. Between-year consistency of migratory choices was high for most storks, but flexible behavior of a few individuals provided an indication of the potential for plastic responses. The coexistence of different migratory strategies in wood storks may be an adaptation to high spatio-temporal heterogeneity and unpredictability of resource availability within their range. Partial migration has been increasingly recognized as a widespread form of migration across taxa, if not the most common (Chapman et al. 2011). Our findings contribute to our general understanding of how bird migration patterns relate to patterns of resource variation in different ecosystems by providing the first individual-based description of migration patterns in a subtropical wading bird.

Our analysis of migratory strategies at the individual level revealed three strategies in the wood stork population: consistent migration, consistent residency, and an intermediate, flexible behavior of facultative migration (Figure 3). Individuals adopting different migratory strategies also differed in their collective seasonal distribution. The distribution of migrants was widely dispersed across the Southeast in the summer and densely concentrated in south Florida in the winter (see overall distribution in Figure 4). This is consistent with previous literature on wood stork seasonal movements (Kahl 1964, Coulter et al. 1999). Migrants likely relocate to south Florida to exploit the winter pulse of food availability in the Everglades as rains cease, pools are isolated and reduced in extent and depth, and fishes are more concentrated and available, and then move north when the rains start, dispersing prey (Kahl 1964, Kushlan 1986). Migration to southern Florida in the winter may also be driven by reduced prey availability in the northern part of the range because of cold temperatures (Frederick and Loftus 1993).

It is unclear whether all migrant storks that spend the winter in south Florida also attempt to nest there. The bimodal distribution we observed for departure dates in the spring (Figure 7A) might result from the fact that some of the migrants leave the winter grounds in south Florida early to go breed elsewhere. An alternative explanation is that among migrant wood storks that attempt to nest in south Florida, those that fail go back to their summer ranges before those that are successful and stay longer to care for their offspring. The existence of different migratory strategies within the population, and particularly of facultative migrants, also suggests that wood storks may behave as “comparison shoppers” when selecting general areas for nesting on any given year. Variable migratory patterns may be associated with variable choices of nesting locations as well, based on a relative comparison of conditions in different parts of the population range. The routes followed by migrant storks varied between seasons, possibly as a response to seasonal variation of thermal air currents which may determine least-cost migratory paths for soaring birds (Kahl 1964, Bohrer et al. 2012, Vansteelant et al. 2017; Figure 7B).

The degree of seasonal range fidelity we observed for migrants suggested that storks tended to repeatedly use the same areas across years, both in winter and summer. For comparison, values of home range overlap corresponding to highest year-to-year breeding site fidelity for wild turkeys (*Meleagris gallopavo*; Badyaev and Faust 1996), on one hand, and capercaillie (*Tetrao urogallus*; *Storch 1997)* and Egyptian vultures (*Neophron percnopterus*; López-López et al. 2014), on the other, are smaller or comparable, respectively, to those we found for wood stork ranges in both seasons. We found that home range size can vary widely for the same individual in different years, possibly according to the degree of dispersion of food resources (Ford 1983, Zabel et al. 1995, Schradin et al. 2010). Consequently, overlap between ranges in different years was rarely exact, but storks tended to return to the same general area equally in summer and winter. Range fidelity may be a critical adaptation to achieve reliable access to resources (Switzer 1993, Vergara et al. 2006), but it might entail susceptibility to changes in habitat quality, which may lead birds into an ecological trap if they remain faithful to areas that were formerly suitable but deteriorated (Schlaepfer et al. 2002, Weldon and Haddad 2005, Lok et al. 2011).

We observed the highest density of year-round residents in southeast Florida – near the northern Everglades and urban coastal areas – and in Jacksonville (Figure 5). Neither of these areas appeared to be intensively used by migrants. The hotspot of resident distribution we observed in the northern Everglades overlaps with the Everglades Agricultural Area (EAA) and Loxahatchee National Wildlife Refuge (i.e. Water Conservation Area (WCA) 1; Figure 5). Water levels are artificially managed throughout the EAA and WCAs through a system of levees and canals according to agricultural schedules and water supply or flood protection needs (Bancroft et al. 2002, Pearlstine et al. 2005). The EAA covers former marsh habitat which was converted to agricultural use starting in the mid-1900s (Pearlstine et al. 2005). In the EAA, whole fields are periodically flooded as part of their crop rotation strategy, often in coincidence with the beginning of the rainy season and rapidly rising water levels in the Everglades (Schueneman et al. 2001, Sizemore 2009, Sizemore and Main 2012). Canals and ditches are periodically drawn down in response to crop needs, and this may provide patches of concentrated fish for foraging wading birds (Pearlstine et al. 2005). The WCAs were impounded in the 1960s with the double purpose of providing water for agricultural and municipal use and flood protection (Light et al. 1989, Light and Dineen 1994). Loxahatchee National Wildlife Refuge is composed of different vegetation communities and characterized by greater micro-topographic relief than other parts of the Everglades, which may provide suitable foraging habitat for storks over a wider temporal range than in situations of uniform topography (Hoffman et al. 1994, Bancroft et al. 2002). The high density of year-round residents we observed in Loxahatchee and the EAA may be ascribed to these features of topography and artificial flooding-and-drying schedules which may result in foraging chances even out of season and out of sync with natural water-level regimes.

We hypothesize that the high concentration of residents near urban areas might be partly linked to the exploitation of supplemental food sources provided deliberately or unintendedly by humans. Resident wood storks in the Jacksonville area were captured at the Jacksonville Zoo. These storks are wild and free-roaming, but regularly receive food supplementation (Bear D., Jacksonville Zoo, personal communication). The high density of storks we observed in Jacksonville might be an artifact of the unequal number of tracked storks at different capture sites, but remarkably most storks captured at the Jacksonville Zoo were consistently resident (7 out of 9). We do not have any direct evidence of food supplementation for storks in southeast Florida, but this is one of the most densely populated urban areas in the Southeast and likely presents several supplementation opportunities. Landfills are a possible source of supplemental food, and there is growing evidence that their use by bird populations, including bald eagles (*Haliaeetus leucocephalus*, Turrin et al. 2015), yellow-legged gulls (*Larus michahellis*, Egunez et al. 2017), and white storks (*Ciconia ciconia*, Gilbert et al. 2016), is increasing in different parts of the world. White ibises (*Eudocimus albus*) have been increasingly observed in the same urban areas of south Florida where we observed the highest concentration of resident storks (Hernandez et al. 2016), and a recent study showed that they heavily rely on artificial food provisioning in urban parks and landfills (Murray et al. 2018). We have anecdotal evidence of wood storks regularly being hand-fed and eating trash in urban environments (Picardi S., personal observation), and ongoing studies on the diet of chicks in urban colonies in southeast Florida have revealed consumption of a diversity of human-derived food that may come from landfills and other sources of trash (Evans B., personal communication). Together, these clues lead us to hypothesize that the availability of supplemental food sources of an artificial nature might be playing a role in determining the distribution of resident storks.

The wood stork population is facing environmental change pressures in many regards, from alterations of the natural hydrological dynamics in the Everglades (Kushlan 1987, Sklar et al. 2001, 2005) to increasing urbanization (Hefner and Brown 1984, Reynolds 2001, Terando et al. 2014), to which the population might respond in the long run by altering migratory patterns. This is an increasingly documented phenomenon in bird populations in response to various drivers, including climate change, changes in resource phenology, and supplemental feeding (Cotton 2003, Visser et al. 2009, Satterfield et al. 2018). Changes in migratory patterns might be expected both through adaptation and behavioral plasticity (Pulido 2007, Ghalambor et al. 2007, Charmantier and Gienapp 2013). Our analysis on consistency of individual migration choices across years highlighted that most of the population (87% of individuals among those monitored over several years) showed highly consistent yearly migratory choices (Figure 3). Notably, the inference we can draw from our data in this sense is limited by the fact that individuals were tracked for only a few years each, if more than one. However, a small proportion of individuals (13% of those monitored over several years) showed some degree of plasticity, making different migratory choices in different years and behaving as facultative migrants (Figure 3). Thus, storks seem to be able to adjust their migratory strategies within the course of a lifetime, implying some potential for plastic changes of migratory behavior at the population level.

Our results provide the first quantitative description of migration patterns in wood storks in the southeastern U.S. and formally establish the status of the population as partially migratory. Partial migration is likely an important adaptation to the exploitation of unpredictable and spatio-temporally heterogeneous resources. The different distribution patterns we observed for migrant and resident individuals suggest that differences in migratory behavior might correspond to different strategies of resource use, whose coexistence may benefit the population by buffering stochasticity of resource availability. While the distribution of migrants reflected seasonal patterns of resource availability across the range, the distribution of residents might be linked to the use of resources made artificially available regardless of seasonality. Consistency of migratory choice was generally high from year to year, but some individuals showed the capability to make different, likely opportunistic migratory choices in different years. Our findings provide an example of partial and facultative migration in a population that relies on resources that are both highly heterogeneous and unpredictable at broad spatio-temporal scales.

Understanding the adaptive significance of partial migration requires a comprehensive assessment of how species inhabiting different ecosystems differ in their migration patterns. By looking at which populations exhibit partial migration or not, researchers can comparatively assess which characteristics of environmental variability lead to its emergence. Studies on multiple species across avian orders have highlighted that partial migration is associated with environments where resource distribution is unstable and highly variable between years (Chan 2001, Jahn et al. 2012). Our study is one of the first to describe and quantify partial migration in a subtropical wading bird. This highlights an important gap in the literature, because partial migration is thought to be associated with heterogeneous and unpredictable environments, of which wetlands are a prime example. Thus, wading birds are good model species to evaluate predictions on partial migration in relation to unpredictable resources. For the same reason, most comparative studies of ecological drivers of avian partial migration have focused on the Australian continent, whose trademark is high climatic variability and unpredictability (Chan 2001). In agreement with previous literature, our findings exemplify that in a highly heterogeneous and unpredictable context, where the availability of key resources varies substantially between years according to variations in rainfall patterns, a combination of partial and facultative migration may be advantageous. Partial migration may buffer between-year stochasticity in survival or reproduction, if the conditions that promote fitness of migrants are different than those of residents. Concurrently, the behavioral flexibility of facultative migrants may work as a reservoir of plasticity, improving population responses to year-to-year variation and allowing rapid change of migratory patterns in response to environmental change.

Future research could focus on comparing migratory patterns across wading bird species to gain deeper understanding of the broad ecological drivers of partial migration. Two types of comparison might yield the most insight: first, comparing migratory patterns in different populations of the same species inhabiting regions subject to different wetland dynamics. Second, comparing migratory patterns of sympatric wading bird species, that despite inhabiting the same watershed may differ in their resource acquisition mechanisms (e.g., searchers vs. exploiters, Gawlik 2002) and thus be more or less sensitive to hydrological fluctuations and respond differently to year-to-year variability.

## Acknowledgments

We thank B.J. Smith for providing helpful comments on earlier versions of this manuscript.

## Author contributions

S.P., P.F., and M.B. conceived the idea and supervised research. R.B. collected the data. S.P. analyzed the data and wrote the paper. All authors contributed to revisions.

## Appendices

### A) Sensitivity analysis of model constraint values

We enforced a spatial and a temporal constraint in the migrant model of Net Squared Displacement by specifying the minimum value of two model parameters, the distance between migratory ranges and the time spent on the second range. The spatial constraint is the minimum distance an individual had to move to be considered a migrant. In our models, we used a value of 260 km. The temporal constraint is the minimum time an individual had to spend in the second range to be considered a migrant. In our models, we used a value of 60 days. Spatial and temporal constraints were applied simultaneously in our models. To ensure that the classification was not affected by the chosen values, we performed a sensitivity analysis on parameter values. We identified a biologically relevant range of values for each constraining parameter. We compared classifications of individual-years resulting from using different values within this range for each of the two parameters, while keeping the other fixed (at 260 km or 60 days). For the spatial constraint, we considered values between 140 km and 500 km, by increments of 30 km. We placed the lower bound of the range at 140 km because wood storks have been observed to travel as far as 130 km from their breeding colony to foraging grounds on a daily scale. Therefore, any value lower than 140 km did not make sense for targeting migratory movements. We chose 500 km as an upper bound because that was approximately the average migration distance we identified during visual inspection of the data. For the temporal constraint, we considered values between 30 and 90 days, by increments of 10 days. We chose 30 days as a lower bound because we found that excursions lasting as long as several weeks were not uncommon during visual inspection of the data. Unlike migrations, excursions lack repeatability and are often at a shorter spatial scale than migrations. We chose 90 days as an upper bound because that was approximately the average time we observed wood storks to spend in the summer range during visual inspection of the data (while they would generally spend longer in the winter range).

Results of the classification corresponding to the use of each possible value of spatial and temporal threshold are reported in Table 1 and Table 2, respectively. For the purpose of this analysis and for simplicity, we only compared the classifications of individual-years for which all models converged at the first iteration (127 and 129 individual-years out of 200 in the case of the spatial and temporal constraint, respectively). Out of 127 individual-years, 108 were consistently classified across the entire range of values considered for the spatial parameter. Out of the 19 that were classified inconsistently, 6 were migrants that failed to be recognized as such for values above 410 km, because the spatial scale of their migration was lower than the threshold (but larger than 380 km); 8 were residents that failed to be recognized as such for values below 230 km, because their daily movements sometimes exceeded that spatial scale (but not 260 km). The remaining 5 individuals were correctly classified as migrants for low threshold values (between 260 and 290 km) and erroneously classified as residents for large values (between 320 and 350 km). Therefore, the classification resulted generally robust between values of 230 and 410 km, with only ∼4% of individual-years classified inconsistently within this range. Within this robust range, the classification of the inconsistent ∼4% individual-years was correct for values lower than 290 km. This supports the use of 260 km as the optimal spatial threshold value to use in the models of migratory behavior. Out of 129 individual-years, 127 were consistently classified across the entire range of values considered for the temporal parameter. Out of the remaining 2, one was a migrant that failed to be recognized as such for values greater than 80 days. The last individual-year was one of the controversial cases that continuously performed non-migratory large scale movements and that we manually assigned to the resident category (see Methods). Thus, the classification was highly robust for the entire range of values considered, and unanimous for values between 30 and 80 days. Overall, results of the sensitivity analysis demonstrate the robustness of our approach, while highlighting variability of the spatial scale of migratory movements in wood storks. This underlines the usefulness and appropriateness of combining a spatial constraint with a temporal constraint, as the latter helps resolving distinctions between migratory and non-migratory movements that would be difficult to disentangle using a spatial criterion alone.

**Table 1:** Results of the classification of wood stork individual-years deriving from the use of different values as a spatial threshold for migratory movements. We used values ranging between 140 and 500 km, by increments of 30 km. The temporal threshold was fixed at 60 days.

| **Value (km)** | **Migrants** | **Residents** |
| --- | --- | --- |
| 140 | 71 | 58 |
| 170 | 70 | 59 |
| 200 | 65 | 64 |
| 230 | 63 | 66 |
| 260 | 61 | 68 |
| 290 | 59 | 70 |
| 320 | 59 | 70 |
| 350 | 57 | 72 |
| 380 | 57 | 72 |
| 410 | 54 | 75 |
| 440 | 55 | 74 |
| 470 | 51 | 78 |
| 500 | 50 | 79 |

**Table 2:** Results of the classification of wood stork individual-years deriving from the use of different values as a temporal threshold for migratory movements. We used values ranging between 30 and 90 days, by increments of 10 days. The spatial threshold was fixed at 260 km.

| **Value (days)** | **Migrants** | **Residents** |
| --- | --- | --- |
| 30 | 62 | 67 |
| 40 | 62 | 67 |
| 50 | 62 | 67 |
| 60 | 61 | 68 |
| 70 | 61 | 68 |
| 80 | 61 | 68 |
| 90 | 60 | 69 |

### B) Starting values for model parameters

The method we used for the classification of migratory behavior relies on the estimation of model parameters, which include the the midpoint of the departing migratory movement, the duration of the migratory movement, the permanence time in the arrival range, the distance between seasonal ranges (in the case of the migratory model), and the average NSD of the resident range and the rate of the initial NSD increase (in the case of the resident model). If starting values for these parameters are not specified by the user, functions in the package migrateR will estimate them based on an internal optimization algorithm. Poor correspondence between starting values of model parameters and the data can impede model convergence. When the models are applied to behaviorally heterogeneous data a single set of starting values for model parameters will likely not be sufficient to ensure convergence of all models. Therefore, a new set of starting values for model parameters can be manually specified in a step-wise process, progressively increasing the number of models that converge until they all do. In our study, 120 models out of 200 converged after the first iteration, and all models converged after 21 iterations with a different set of starting values. We report the sets of starting values we used in Table 3.

**Table 3:** Starting parameter values used in our analysis to progressively improve model convergence until full convergence. Sets of values are reported in the order in which we used them.

| **Distance between**<br>**ranges (km)** | **Mean NSD of starting**<br>**range (km<sup>2</sup>)** | **Time to complete ½ to**<br>**¾ of migratory**<br>**movement (days)** | **Failed convergences**<br>**(n)** |
| --- | --- | --- | --- |
| Not specified | Not specified | Not specified | 80 |
| 260 | Not specified | Not specified | 73 |
| 300 | Not specified | Not specified | 67 |
| 400 | Not specified | Not specified | 66 |
| 500 | Not specified | Not specified | 64 |
| 550 | Not specified | Not specified | 63 |
| Not specified | 100 | Not specified | 36 |
| Not specified | 400 | Not specified | 24 |
| Not specified | 900 | Not specified | 18 |
| Not specified | 1600 | Not specified | 15 |
| Not specified | 3600 | Not specified | 13 |
| Not specified | 4900 | Not specified | 10 |
| Not specified | 8100 | Not specified | 8 |
| Not specified | 10000 | Not specified | 7 |
| Not specified | 14400 | Not specified | 6 |
| Not specified | 28900 | Not specified | 5 |
| Not specified | Not specified | 3 | 4 |
| Not specified | Not specified | 12 | 3 |
| Not specified | Not specified | 13 | 2 |
| Not specified | 100000 | Not specified | 1 |
| Not specified | 650000 | Not specified | 0 |

### C) Discarded individual-years

We discarded 12 individual-years from subsequent analyses because the temporal extent or arrangement of their data did not allow us to unequivocally classify them (Figures 8-19).

## Literature Cited

Baber, M. J., D. L. Childers, K. J. Babbitt, and D. H. Anderson (2002). Controls on fish distribution and abundance in temporary wetlands. Canadian Journal of Fisheries and Aquatic Sciences 59:1441–1450.

Badyaev, A. V., and J. D. Faust (1996). Nest Site Fidelity in Female Wild Turkey: Potential Causes and Reproductive Consequences. The Condor 98:589–594.

Bancroft, G. T., D. E. Gawlik, and K. Rutchey (2002). Distribution of Wading Birds Relative to Vegetation and Water Depths in the Northern Everglades of Florida, USA. Waterbirds 25:265–277.

Beerens, J. M. (2008). Heirarchical resource selection and movements of two wading bird species with divergent foraging strategies in the Everglades. [Online.] Available at https://search.proquest.com/docview/304568853/abstract/E7EAD8D8C1094AA7PQ/1.

Beissinger, S. R. (1995). Modeling Extinction in Periodic Environments: Everglades Water Levels and Snail Kite Population Viability. Ecological Applications 5:618–631.

Berthold, P. (2001). Bird Migration: A General Survey. Oxford University Press.

Bohrer, G., D. Brandes, J. T. Mandel, K. L. Bildstein, T. A. Miller, M. Lanzone, T. Katzner, C. Maisonneuve, and J. A. Tremblay (2012). Estimating updraft velocity components over large spatial scales: contrasting migration strategies of golden eagles and turkey vultures. Ecology Letters 15:96–103.

Börger, L., N. Franconi, G. De Michele, A. Gantz, F. Meschi, A. Manica, S. Lovari, and T. Coulson (2006). Effects of sampling regime on the mean and variance of home range size estimates. Journal of Animal Ecology 75:1393–1405.

Borkhataria, R. R., A. Lawrence Bryan, and P. C. Frederick (2013). Movements and Habitat Use by Fledgling Wood Storks (Mycteria Americana) Prior to Dispersal from the Natal Colony. Waterbirds 36:409–417.

Botson, B. A., D. E. Gawlik, and J. C. Trexler (2016). Mechanisms That Generate Resource Pulses in a Fluctuating Wetland. PLOS ONE 11:e0158864.

Bryan, A. L., W. B. Brooks, J. D. Taylor, D. M. Richardson, C. W. Jeske, and I. L. Brisbin (2008). Satellite Tracking Large-scale Movements of Wood Storks Captured in the Gulf Coast Region. Waterbirds 31:35–41.

Bryan, A. L., and M. C. Coulter (1987). Foraging Flight Characteristics of Wood Storks in East-Central Georgia, U.S.A. Colonial Waterbirds 10:157–161.

Bunnefeld, N., L. Börger, B. van Moorter, C. M. Rolandsen, H. Dettki, E. J. Solberg, and G. Ericsson (2011). A model driven approach to quantify migration patterns: individual, regional and yearly differences. Journal of Animal Ecology 80:466–476.

Calenge, C. (2006). The package “adehabitat” for the R software: A tool for the analysis of space and habitat use by animals. Ecological Modelling 197:516–519.

Calenge, C., S. Dray, and M. Royer-Carenzi (2009). The concept of animals’ trajectories from a data analysis perspective. Ecological Informatics 4:34–41.

Chan, K. (2001). Partial migration in Australian landbirds: a review. Emu - Austral Ornithology 101:281–292.

Chapman, B. B., C. Brönmark, J.-Å. Nilsson, and L.-A. Hansson (2011). The ecology and evolution of partial migration. Oikos 120:1764–1775.

Charmantier, A., and P. Gienapp (2013). Climate change and timing of avian breeding and migration: evolutionary versus plastic changes. Evolutionary Applications 7:15–28.

Chick, J. H., C. R. Ruetz, and J. C. Trexler (2004). Spatial scale and abundance patterns of large fish communities in freshwater marshes of the Florida Everglades. Wetlands 24:652–664.

Cotton, P. A. (2003). Avian migration phenology and global climate change. Proceedings of the National Academy of Sciences 100:12219–12222.

Coulter, M. C., J. A. Rodgers, J. C. Ogden, and F. C. Depkin (1999). Wood Stork(Mycteria americana). The Birds of North America:24.

Cox, G. W. (1985). The Evolution of Avian Migration Systems between Temperate and Tropical Regions of the New World. The American Naturalist 126:451–474.

DeAngelis, D. L., J. C. Trexler, C. Cosner, A. Obaza, and F. Jopp (2010). Fish population dynamics in a seasonally varying wetland. Ecological Modelling 221:1131–1137.

DeAngelis, D. L., J. C. Trexler, and W. F. Loftus (2005). Life history trade-offs and community dynamics of small fishes in a seasonally pulsed wetland. Canadian Journal of Fisheries and Aquatic Sciences 62:781–790.

Dingle, H. (1996). Migration: The Biology of Animals on the Move. Oxford University Press.

Dingle, H., and V. A. Drake (2007). What Is Migration? BioScience 57:113–121.

Dukai, B., M. Basille, and D. Bucklin (2016). rpostgisLT: Managing Animal Movement Data with ‘PostGIS’ and R.

Egunez, A., N. Zorrozua, A. Aldalur, A. Herrero, and J. Arizaga (2017). Local use of landfills by a yellow-legged gull population suggests distance-dependent resource exploitation. Journal of Avian Biology 49:jav–01455.

Fletcher, Jr., Robert J., and R. R. Koford (2004). Consequences of rainfall variation for breeding wetland blackbirds. Canadian Journal of Zoology 82:1316–1325.

Ford, R. G. (1983). Home Range in a Patchy Environment: Optimal Foraging Predictions. Integrative and Comparative Biology 23:315–326.

Frederick, P. C., and W. F. Loftus (1993). Responses of Marsh Fishes and Breeding Wading Birds to Low Temperatures: A Possible Behavioral Link between Predator and Prey. Estuaries 16:216.

Frederick, P. C., and J. C. Ogden (1997). Philopatry and Nomadism: Contrasting Long-Term Movement Behavior and Population Dynamics of White Ibises and Wood Storks. Colonial Waterbirds 20:316.

Frederick, P. C., and J. C. Ogden (2001). Pulsed breeding of long-legged wading birds and the importance of infrequent severe drought conditions in the Florida Everglades. Wetlands 21:484–491.

Frederick, P., D. E. Gawlik, J. C. Ogden, M. I. Cook, and M. Lusk (2009). The White Ibis and Wood Stork as indicators for restoration of the everglades ecosystem. Ecological Indicators 9:S83–S95.

Gawlik, D. E. (2002). The Effects of Prey Availability on the Numerical Response of Wading Birds. Ecological Monographs 72:329–346.

Ghalambor, C. K., J. K. McKAY, S. P. Carroll, and D. N. Reznick (2007). Adaptive versus non-adaptive phenotypic plasticity and the potential for contemporary adaptation in new environments. Functional Ecology 21:394–407.

Gilbert, N. I., R. A. Correia, J. P. Silva, C. Pacheco, I. Catry, P. W. Atkinson, J. A. Gill, and A. M. A. Franco (2016). Are white storks addicted to junk food? Impacts of landfill use on the movement and behaviour of resident white storks (Ciconia ciconia) from a partially migratory population. Movement Ecology 4:7.

Haig, S. M., D. W. Mehlman, and L. W. Oring (1998). Avian Movements and Wetland Connectivity in Landscape Conservation. Conservation Biology 12:749–758.

Hefner, J. M., and J. D. Brown (1984). Wetland trends in the Southeastern United States. Wetlands 4:1–11.

Hegemann, A., A. M. Fudickar, and J.-Å. Nilsson (2019). A physiological perspective on the ecology and evolution of partial migration. Journal of Ornithology. https://doi.org/10.1007/s10336-019-01648-9

Hernandez, S. M., C. N. Welch, V. E. Peters, E. K. Lipp, S. Curry, M. J. Yabsley, S. Sanchez, A. Presotto, P. Gerner-Smidt, K. B. Hise, E. Hammond, et al. (2016). Urbanized White Ibises (Eudocimus albus) as Carriers of Salmonella enterica of Significance to Public Health and Wildlife. PLOS ONE 11:e0164402.

Herring, G. (2008). Constraints of landscape level prey availability on physiological condition and productivity of Great Egrets and White Ibises in the Florida Everglades. Florida Atlantic University.

Hoffman, W., G. T. Bancroft, and R. J. Sawicki (1994). Foraging habitat of wading birds in the Water Conservation Areas of the Everglades. Everglades: the ecosystem and its restoration:585–614.

Hylton, R. A. (2004). Survival, movement patterns, and habitat use of juvenile Wood Storks, Mycteria americana. [Online.] Available at http://purl.fcla.edu/fcla/etd/UFE0007007.

Jahn, A. E., S. P. Bravo, V. R. Cueto, D. J. Levey, and M. V. Morales (2012). Patterns of partial avian migration in northern and southern temperate latitudes of the New World. Emu - Austral Ornithology 112:17–22.

Junk, W. J. (1993). Wetlands of tropical South America. In Wetlands of the world: Inventory, ecology and management Volume I: Africa, Australia, Canada and Greenland, Mediterranean, Mexico, Papua New Guinea, South Asia, Tropical South America, United States (D. F. Whigham, D. Dykyjová and S. Hejný, Editors). Springer Netherlands, Dordrecht, pp. 679–739.

Kahl, M. P. (1964). Food Ecology of the Wood Stork (Mycteria americana) in Florida. Ecological Monographs 34:97–117.

Kareiva, P. M., and N. Shigesada (1983). Analyzing insect movement as a correlated random walk. Oecologia 56:234–238.

Kingsford, R. T., D. A. Roshier, and J. L. Porter (2010). Australian waterbirds – time and space travellers in dynamic desert landscapes. Marine and Freshwater Research 61:875.

Kushlan, J. A. (1981). Resource Use Strategies of Wading Birds. The Wilson Bulletin 93:145– 163.

Kushlan, J. A. (1986). Responses of Wading Birds to Seasonally Fluctuating Water Levels: Strategies and Their Limits. Colonial Waterbirds 9:155–162.

Kushlan, J. A. (1987). External threats and internal management: the hydrologic regulation of the Everglades, Florida, USA. Environmental Management 11:109–119.

Light, S. S., and J. W. Dineen (1994). Water control in the Everglades: a historical perspective. Everglades: The ecosystem and its restoration 5:47–84.

Light, S. S., J. R. Wodraska, and S. Joe (1989). The Southern Everglades. National Forum; Baton Rouge, La. 69:11–14.

Loftus, W. F., and A.-M. Eklund (1994). Long-term Dynamics of an Everglades Small-fish Assemblage. In Everglades: The Ecosystem and Its Restoration. CRC Press.

Lok, T., O. Overdijk, J. M. Tinbergen, and T. Piersma (2011). The paradox of spoonbill migration: most birds travel to where survival rates are lowest. Animal Behaviour 82:837–844.

López-López, P., C. García-Ripollés, and V. Urios (2014). Food predictability determines space use of endangered vultures: implications for management of supplementary feeding. Ecological Applications 24:938–949.

Mckilligan, N. G., D. S. Reimer, D. H. C. Seton, D. H. C. Davidson, and J. T. Willows (1993). Survival and Seasonal Movements of the Cattle Egret in Eastern Australia. Emu 93:79– 87.

Melvin, S. L., D. E. Gawlik, and T. Scharff (1999). Long-Term Movement Patterns for Seven Species of Wading Birds. Waterbirds: The International Journal of Waterbird Biology 22:411–416.

Murray, M. H., A. D. Kidd, S. E. Curry, J. Hepinstall-Cymerman, M. J. Yabsley, H. C. Adams, T. Ellison, C. N. Welch, and S. M. Hernandez (2018). From wetland specialist to hand-fed generalist: shifts in diet and condition with provisioning for a recently urbanized wading bird. Phil. Trans. R. Soc. B 373:20170100.

Nakagawa, S., and H. Schielzeth (2013). A general and simple method for obtaining R2 from generalized linear mixed-effects models. Methods in Ecology and Evolution 4:133–142.

Newton, I. (2012). Obligate and facultative migration in birds: ecological aspects. Journal of Ornithology 153:171–180.

Niemuth, N. D., M. E. Estey, R. E. Reynolds, C. R. Loesch, and W. A. Meeks (2006). Use of wetlands by spring-migrant shorebirds in agricultural landscapes of North Dakota’s Drift Prairie. Wetlands 26:30–39.

Niemuth, N. D., and J. W. Solberg (2003). Response of Waterbirds to Number of Wetlands in the Prairie Pothole Region of North Dakota, U.S.A. Waterbirds 26:233–238.

Ogden, J. C. (1986). The Wood Stork. In Audubon Wildlife Report 1985. Di Silvestro, R. L. The National Audubon Society, New York, New York, USA, pp. 458–471.

Ogden, J. C., J. A. Kushlan, and J. T. Tilmant (1976). Prey Selectivity by the Wood Stork. The Condor 78:324–330.

Ogden, J. C., J. A. Kushlan, and J. T. Tilmant (1978). The food habits and nesting success of Wood Storks in Everglades National Park 1974. Department of the Interior, National Park Service.

Pearlstine, E. V., M. L. Casler, and F. J. Mazzotti (2005). A checklist of birds of the Everglades Agricultural Area. Florida Scientist:84–96.

Picardi, S., R. R. Borkhataria, A. L. B. Jr, P. C. Frederick, and M. Basille (2018). GPS Telemetry Reveals Occasional Dispersal of Wood Storks from the Southeastern US to Mexico. 7.

Poiani, A. (2006). Effects of Floods on Distribution and Reproduction of Aquatic Birds. In Advances in Ecological Research. Academic Press, pp. 63–83.

Pulido, F. (2007). Phenotypic changes in spring arrival:: evolution, phenotypic plasticity, effects of weather and condition. Climate Research 35:5–23.

Reynolds, J. E. (2001). Urbanization and land use change in Florida and the South. Current Issues Associated with Land Values and Land Use Planning Proceedings of a Regional Workshop. p. 28.

Roshier, D. A., V. A. J. Doerr, and E. D. Doerr (2008). Animal movement in dynamic landscapes: interaction between behavioural strategies and resource distributions. Oecologia 156:465–477.

Ruetz, C. R., J. C. Trexler, F. Jordan, W. F. Loftus, and S. A. Perry (2005). Population dynamics of wetland fishes: spatio-temporal patterns synchronized by hydrological disturbance? Journal of Animal Ecology 74:322–332.

Satterfield, D. A., P. P. Marra, T. S. Sillett, and S. Altizer (2018). Responses of migratory species and their pathogens to supplemental feeding. Phil. Trans. R. Soc. B 373:20170094.

Schlaepfer, M. A., M. C. Runge, and P. W. Sherman (2002). Ecological and evolutionary traps. Trends in Ecology & Evolution 17:474–480.

Schradin, C., G. Schmohl, H. G. Rödel, I. Schoepf, S. M. Treffler, J. Brenner, M. Bleeker, M. Schubert, B. König, and N. Pillay (2010). Female home range size is regulated by resource distribution and intraspecific competition: a long-term field study. Animal Behaviour 79:195–203.

Schueneman, T. J., C. Rainbolt, and R. Gilbert (2001). Rice in the crop rotation. University of Florida Cooperative Extension Service, Institute of Food and Agricultural Sciences, EDIS.

Sekercioglu, C. H. (2010). Partial migration in tropical birds: the frontier of movement ecology. Journal of Animal Ecology 79:933–936.

Sergio, F., J. Blas, L. López, A. Tanferna, R. Díaz-Delgado, J. A. Donázar, and F. Hiraldo (2011). Coping with uncertainty: breeding adjustments to an unpredictable environment in an opportunistic raptor. Oecologia 166:79–90.

Sizemore, G. (2009). Foraging quality of flooded agricultural fields within the Everglades Agricultural Area for wading birds (Ciconiiformes).

Sizemore, G. C., and M. B. Main (2012). Quality of Flooded Rice and Fallow Fields as Foraging Habitat for Little Blue Herons and Great Egrets in the Everglades Agricultural Area, USA. Waterbirds 35:381–393.

Sklar, F. H., M. J. Chimney, S. Newman, P. McCormick, D. Gawlik, S. Miao, C. McVoy, W. Said, J. Newman, C. Coronado, G. Crozier, et al. (2005). The ecological–societal underpinnings of Everglades restoration. Frontiers in Ecology and the Environment 3:161–169.

Sklar, F., C. McVoy, R. VanZee, D. Gawlik, K. Tarboton, D. Rudnick, S. Miao, and T. Armentano (2001). The Effects of Altered Hydrology on the Ecology of the Everglades. In The Everglades, Florida Bay, and Coral Reefs of the Florida Keys. CRC Press.

Snodgrass, J. W., A. L. Bryan, R. F. Lide, and G. M. Smith (1996). Factors affecting the occurrence and structure of fish assemblages in isolated wetlands of the upper coastal plain, U.S.A. 53:12.

Spitz, D. B., M. Hebblewhite, and T. R. Stephenson (2017). ‘MigrateR’: extending model-driven methods for classifying and quantifying animal movement behavior. Ecography 40:788– 799.

Storch, I. (1997). Male territoriality, female range use, and spatial organisation of capercaillie *Tetrao urogallus* leks. Wildlife Biology 3:149–161.

Streich, W. J., H. Litzbarski, B. Ludwig, and S. Ludwig (2006). What triggers facultative winter migration of Great Bustard (Otis tarda) in Central Europe? European Journal of Wildlife Research 52:48–53.

Switzer, P. V. (1993). Site fidelity in predictable and unpredictable habitats. Evolutionary Ecology 7:533–555.

Team, R. C. (2018). R: A Language and Environment for Statistical Computing, R Foundation for Statistical Computing, Austria, 2015. ISBN3-900051-07-0: URL http://www.R-project.org.

Terando, A. J., J. Costanza, C. Belyea, R. R. Dunn, A. McKerrow, and J. A. Collazo (2014). The Southern Megalopolis: Using the Past to Predict the Future of Urban Sprawl in the Southeast U.S. PLOS ONE 9:e102261.

Turrin, C., B. D. Watts, and E. K. Mojica (2015). Landfill Use by Bald Eagles in the Chesapeake Bay Region. Journal of Raptor Research 49:239–249.

Van Moorter, B., N. Bunnefeld, M. Panzacchi, C. M. Rolandsen, E. J. Solberg, and B.-E. Sæther (2013). Understanding scales of movement: animals ride waves and ripples of environmental change. Journal of Animal Ecology 82:770–780.

Vansteelant, W. M. G., J. Shamoun Baranes, W. van Manen, J. van Diermen, and W. Bouten (2017). Seasonal detours by soaring migrants shaped by wind regimes along the East Atlantic Flyway. Journal of Animal Ecology 86:179–191.

Vergara, P., J. I. Aguirre, J. A. Fargallo, and J. A. Dávila (2006). Nest-site fidelity and breeding success in White Stork Ciconia ciconia. Ibis 148:672–677.

Visser, M. E., A. C. Perdeck, J. H. V. Balen, and C. Both (2009). Climate change leads to decreasing bird migration distances. Global Change Biology 15:1859–1865.

Weldon, A. J., and N. M. Haddad (2005). The Effects of Patch Shape on Indigo Buntings: Evidence for an Ecological Trap. Ecology 86:1422–1431.

Weller, M. W. (1999). Wetland Birds: Habitat Resources and Conservation Implications. Cambridge University Press.

Zabel, C. J., K. McKelvey, and J. P. Ward Jr. (1995). Influence of primary prey on home-range size and habitat-use patterns of northern spotted owls (Strix occidentalis caurina). Canadian Journal of Zoology 73:433–439.

